# Functional-based parcellation of the mouse prefrontal cortex for network perturbation analysis

**DOI:** 10.1101/2024.03.25.586664

**Authors:** Iurii Savvateev, Christina Grimm, Marija Markicevic, Joanes Grandjean, Nicole Wenderoth, Rafael Polania, Valerio Zerbi

## Abstract

The prefrontal cortex (PFC) is a region of the brain involved in higher-order cognitive processes such as attention, emotional regulation, and social behavior, making it a hotspot for an ongoing clinical and fundamental research. Importantly, its functionality is intricately interconnected within a wide array of functional networks encompassing multiple other brain areas. However, the delineation of distinct subdivisions within the mouse PFC and their contributions to the broader brain network function remain topics of ongoing debate. In the current study, we used resting-state functional magnetic resonance imaging (rsfMRI) from a large cohort of wildtype animals to derive the functional-connectivity (FC) based parcellation of the mouse PFC with voxel resolution. Our findings indicate the presence of FC-based clusters that deviate from the established anatomical subdivisions within the cingulate and prelimbic areas, while they align in infralimbic and orbital cortices. To further underscore the association of these FC-based clusters with distinct functional networks, we performed network-specific perturbations using chemogenetics in the identified clusters in dorsal PFC and monitor the elicited effects with fMRI (chemo-fMRI). Our recordings revealed that FC perturbations were observed only within the functional networks linked to the targeted clusters and did not spread to neighbouring anatomical areas or functional clusters. We propose that FC-based parcellation is a valuable approach for tracking the impact of external activations and confirming the precise site of activation.

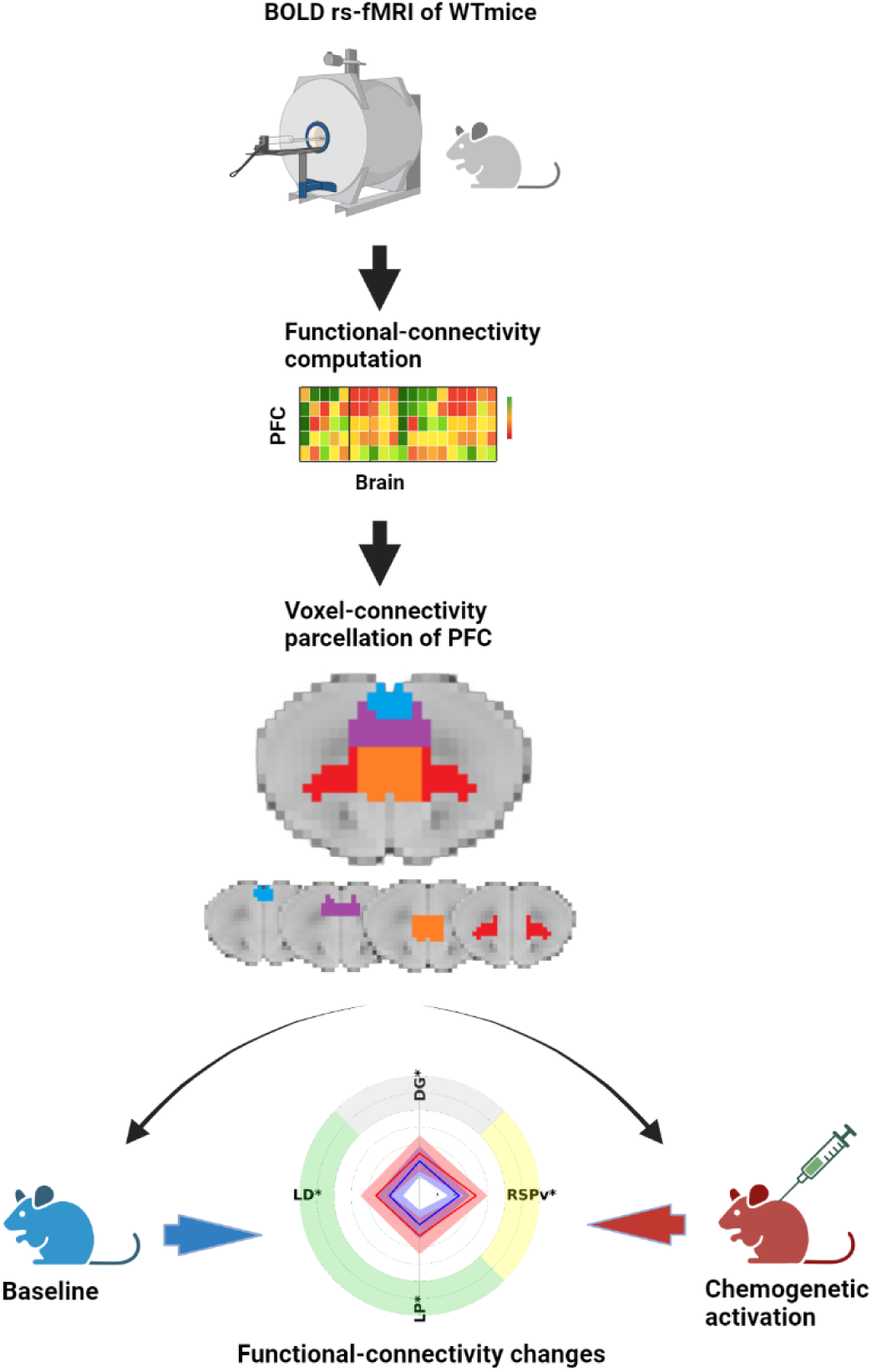

## Introduction

The prefrontal cortex (PFC) is a central hub in the brain that integrates sensory information, emotions, and previous experience to orchestrate the design and execution of most suitable responses (Le Merre, Ahrlund-Richter, & Carlen, 2021). Early research on the PFC in macaques ignited an interest in the area (Fuster & Alexander, 1971), which has since grown exponentially and is currently being investigated in a broad range of species (Laubach, Amarante, Swanson, & White, 2018). Among them, the ability to study these circuits in mouse models, which allow for much more direct neuronal, pharmacological, and genetic manipulation of the system, would be extremely beneficial for the development and validation of novel therapies (Carlén, 2017; Laubach et al., 2018). PFC circuits in rodents have long been a source of controversy, with disagreements centred on (i) their location, subdivisions and functions (Le Merre et al., 2021) as well as on (ii) their homology with human areas (Bicks, Koike, Akbarian, & Morishita, 2015). The precise location of the border between primary prelimbic, infralimbic areas and anterior cingulate is particularly contentious (Laubach et al., 2018; Le Merre et al., 2021) and is periodically revised in anatomical atlases (Van de Werd & Uylings, 2014). Moreover, homologies between mouse PFC and human areas remain mostly speculative as differences in the gross anatomy make it difficult to establish parallels in the organization of the PFC between the human and rodent brains (Bicks et al., 2015). Hence, new ways to define functional regions and circuits in mice are required.

Common approaches in rodents involves the analysis of cytoarchitecture, anatomical connectivity, and gene expression profiles (Paxinos & Franklin, 2019; Wang et al., 2020). While such an anatomy-based parcellation can inform researchers about the basic organization of the PFC, it is not exempt from a set of challenges. This includes nomenclature-related discrepancies across species as a result of (i) using different name schemes within rodent studies (Laubach et al., 2018; Paxinos & Franklin, 2019; Wang et al., 2020) and (ii) inconsistent boundaries amongst homologous regions across species (van Heukelum et al., 2020). Consequently, rather than focusing on common structures, it is advantageous to research PFC function in respect to its various long-range inputs and outputs from and to cortical and subcortical locations (Anastasiades & Carter, 2021; Wilson, Gaffan, Browning, & Baxter, 2010), i.e. at the level of whole-brain functional networks (Zerbi, Grandjean, Rudin, & Wenderoth, 2015).

Whole-brain imaging techniques such as functional Magnetic Resonance Imaging (fMRI) allow researchers to derive functional connectivity (i.e. FC) based parcellation relying on orchestrated neural activity, thus, partially circumventing the challenges associated with an anatomy-based parcellation. The process of FC-based parcellation consists of two principal steps. First, by computing pairwise correlations of the signals from different voxels/brain areas the corresponding functional connectivity (FC) is estimated. Second, the generated correlational matrix is further clustered using, for instance, methods from machine learning or graph theory (Arslan et al., 2018). Over the last years FC based parcellations using both resting-state and task-based fMRI were applied in humans to build whole-brain and regional parcellations (Arslan et al., 2018; James, Hazaroglu, & Bush, 2016; Moghimi et al., 2022). Importantly, the interspecies comparison of FC-based parcellations allowed to refine the concept of homologous brain areas. For instance, the comparison of monkeys and human visual cortices revealed that high-order visual processing areas in monkeys (e.g. V3A) are functionally related to regions located differently as predicted by anatomical correspondence (Mantini et al., 2012).

Rodent fMRI with current state-of-the-art protocols provide remarkable SNR capabilities at high temporal and spatial resolutions that are compatible with recent advancements in human fMRI (Coletta et al., 2020; Grandjean, Zerbi, Balsters, Wenderoth, & Rudin, 2017; Mantini et al., 2012). Combined with the circuitry perturbations done by optogenetics or chemogenetics permitting the stimulation of specific cell populations, rodent fMRI allows scientists to discern the functional roles of the targeted cell in a whole-brain functional connectome (Mandino et al., 2019; Markicevic et al., 2020). Here, we conducted a study with two main objectives. First, we aimed to provide a functional parcellation of the mouse PFC at voxel resolution leveraging a database of 100 resting-state fMRI scans obtained during light anesthesia in wildtype C57BL6/j mice (Zerbi et al., 2021). Our results demonstrate that functional connectivity (FC) based clustering can capture the mesoscopic-level PFC subdivisions previously reported by axonal tracing studies (Harris et al., 2019; Le Merre et al., 2021). However, we also described discrepancies between the spatial arrangements of functional clusters and the anatomically defined borders within the mPFC (Laubach et al., 2018). Second, we leveraged our FC-based parcellation to manipulate the excitation: inhibition balance (E:I) in a network-specific manner. Our findings revealed functional connectivity changes that were restricted to the functional clusters tested and did not extend to neighbouring but functionally isolated clusters. These results establish a functional segregation of the mPFC in mice and opens new avenues for experimental perturbational protocols based on network mapping.

## Results

### FC based clustering of resting-state fMRI data reveals a subdivision of PFC in accordance with the Allen Parcellation atlas

In this study we aimed to segment PFC using FC data. Firstly, we extracted blood-oxygen-level-dependent (BOLD) time series in each voxel of the brain and calculated Pearson’s correlations between 995 voxels assigned to PFC based on the Allen Mouse Brain Common Coordinate Framework version 3 (CCFv3) (Fig 1A)(Wang et al., 2020) and 16931 voxels from the rest of the brain. The correlational matrices (995x16931) were computed for a subset of 60 mice, randomly chosen from a database of 100 resting-state recordings of C57BL/6J adult mice (Zerbi et al., 2021). The resulting matrix (995x16931x60) was averaged across animals and the mean correlational matrix (995x16931) (Fig 1B) was clustered using k-means. The assessment of silhouette scores and inertia demonstrated that 4 was the best cluster solution, leading to the highest silhouette score and lying at the elbow of the inertia plot (Fig 1 C). The revealed four clusters showed a separation along the rostro-caudal axis, matching qualitatively the demarcation used in CCFv3. Low-dimensional representation of the correlational matrix further revealed a strong similarity in the internal structure of the data which mimics the anatomical structure of PFC (both 3D and 2D, Fig 1 D).

**Figure 1.**
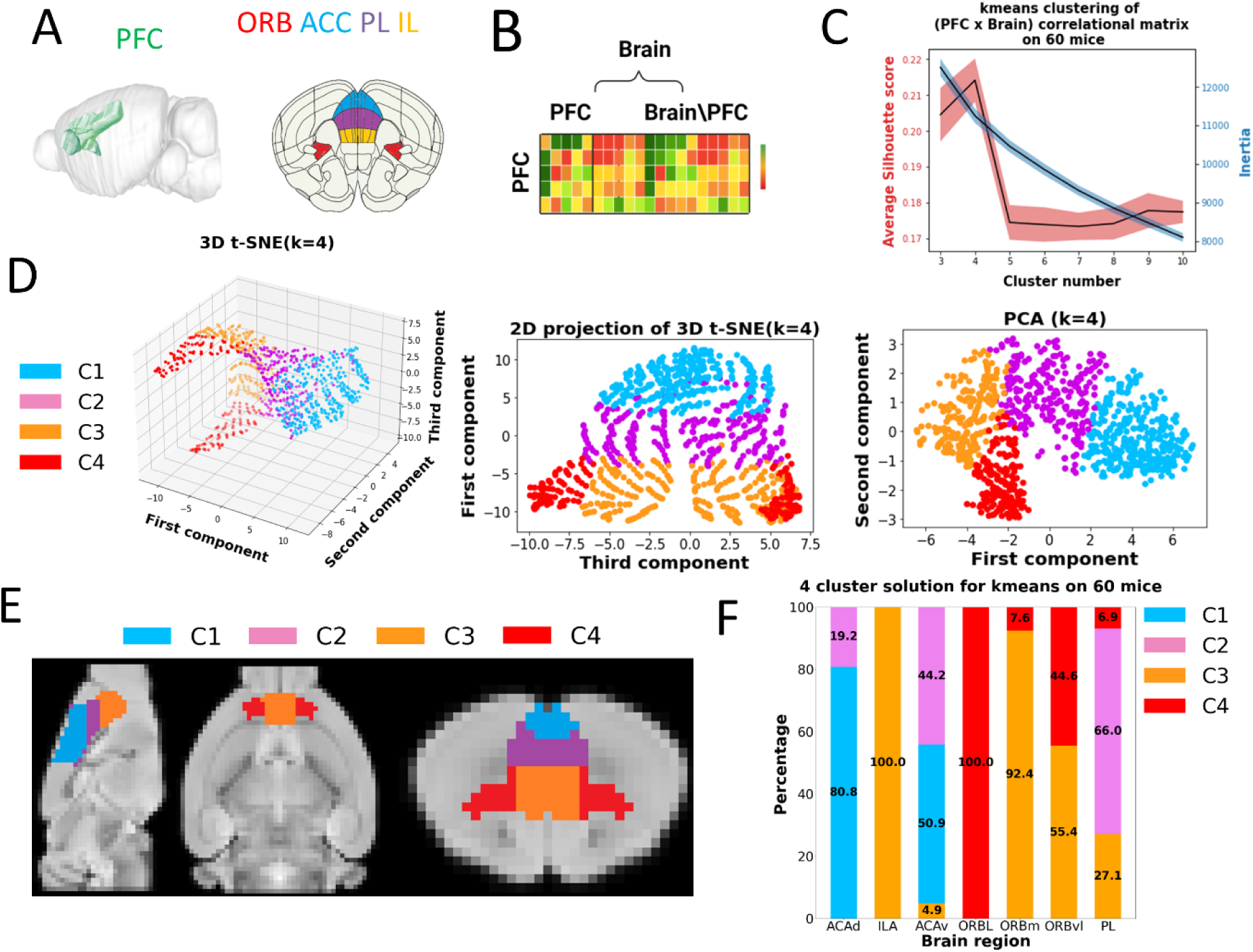
PFC clustering based on the whole-brain functional connectivity. **A:** Visualization of PFC and its subregions based on Allen Mouse Brain Common Coordinate Framework version 3 (CCFv3). **B:** Scheme of the correlation matrix used to derive PFC subdivisions via kmeans. **C:** Average silhouette and inertia scores for various cluster solutions using 60 randomly chosen mice. Black line represents the mean values across 200 iterations. Shaded area features the standard deviation. **D:** Low dimensional representation of the correlation matrix through 3D t-SNE and PCA for 4-cluster solution. Colors represent the clusters derived by k-means clustering on the complete (not-embedded) correlation matrix. **E:** Visualization of the anatomy of individual clusters along sagittal, transverse and coronal sections. **F:** Prevalence of individual clusters based on 4-cluster solution within anatomically defined (CCFv3) regions. Percentage values are depicted only for the regions with a contribution above 3%.

To ensure the reproducibility of the identified cluster solution, we repeated the clustering using different data sizes and clustering techniques (Fig S1 and Fig S2). The stability of the 4-cluster solution was corroborated by the invariance of clusters borders under the usage of different data sizes (see examples in Fig S1 B,D,F). The notion that the 4-cluster solution is an optimal choice regardless of data size emphasizes the homogeneity of the investigated data and the stability of the obtained solution (Fig S1 A,C,E). To ensure that the shapes of the clusters are method-independent we further perform Gaussian-Mixture-Models (GMM) and Hierarchical clustering on the complete 100 mice dataset (Fig S2).

Next, we calculated the contribution (%) of the four clusters to the seven PFC sub-region defined by CCFv3. We observed that five sub-regions identified by CCFv3 were mostly integrated in one cluster: ACA(d)-C1, PL-C2, IL-C3. ORB(l)-C4, ORB(m)-C3. The ORB(vl) and ACA(v) were instead shared by two clusters: C3/C4 and C1/C2 respectively (Fig 1 F).

The higher prevalence of C4 in ORB(l) (100%) and ORB(vl) (44.6%) compared to ORBm (7.6%) is in line with the connection-density based PFC segregation: ORB(l) and ORB(vl) were assigned to vlPFC, whereas ORBm was linked to vmPFC (Le Merre et al., 2021). Moreover, ORB(l)/ORB(vl) and ORB(m) were assigned to different functional networks, named "medial networks"(Zingg et al., 2014). ORB(l) and ORB(vl) were linked to the subnetwork performing higher level processing of somatosensory, auditory, and visual data and connecting processed sensory information with motor actions. At the same time ORB(m), was linked to the network together with PL/IL and associated with the processing of spatial information transmitted from the dorsal subiculum (SUBd) via ventral retrosplenial area (RSPv) and ACAv(Fanselow & Dong, 2010; Zingg et al., 2014). The functional coupling of ORB(m) with ILA is further corroborated by the dominance of the same functional cluster C3 in both: 100% for ILA and 92.4% (Fig 1F, Table S1).

We found contributions from PL-dominant (66%) cluster C2 in both parts of ACA: ACAv and ACAd. This is in accordance with the existing anatomy-based overlaps of ACA and PL. Specifically, ACAd and ACAv have a common anatomical border with PL (van Heukelum et al., 2020) which exact location is not consistent across the studies (Le Merre et al., 2021) suggesting ACA and PL will likely share common functional and connectivity features. By comparing the prevalence of C2 in ACA(v) and ACA(d): 44.2% and 19.2% respectively we could suggest that most of the functional overlap occurs in the ventral part of ACAv. The greater functional overlap between ACA(v) and PL is also consistent with the above discussed involvement of mostly ventral ACA in relaying spatial information from SUBd to PL/IL(Zingg et al., 2014).

### FC based clustering captures high-order PFC subdivision

We then proceeded to test the influence of the short-range (within PFC) and long-range (between PFC and the rest of the brain) functional connectivity on our clustering. Based on the reported extensive interconnectivity within PFC (Harris et al., 2019; Le Merre et al., 2021; Zingg et al., 2014) we hypothesized that local FC may play a prevalent role when a high-order PFC subdivision takes place. To do that, we used the same FC-based parcellation pipeline as described above for the whole-brain FC (PFC vs Brain) but using either [PFC vs PFC] (995x995) or [PFC vs Brain without PFC] (995x 15936) correlational matrices, similarly as previously done in (Cha, Jo, Gibson, & Lee, 2017).

We observed that 4-cluster was the optimal solution for both matrices (Fig 2C and Fig S3) and that it was invariant to changes in the amount of the examined data (Fig S3 and Fig. S4).

**Figure 2.**
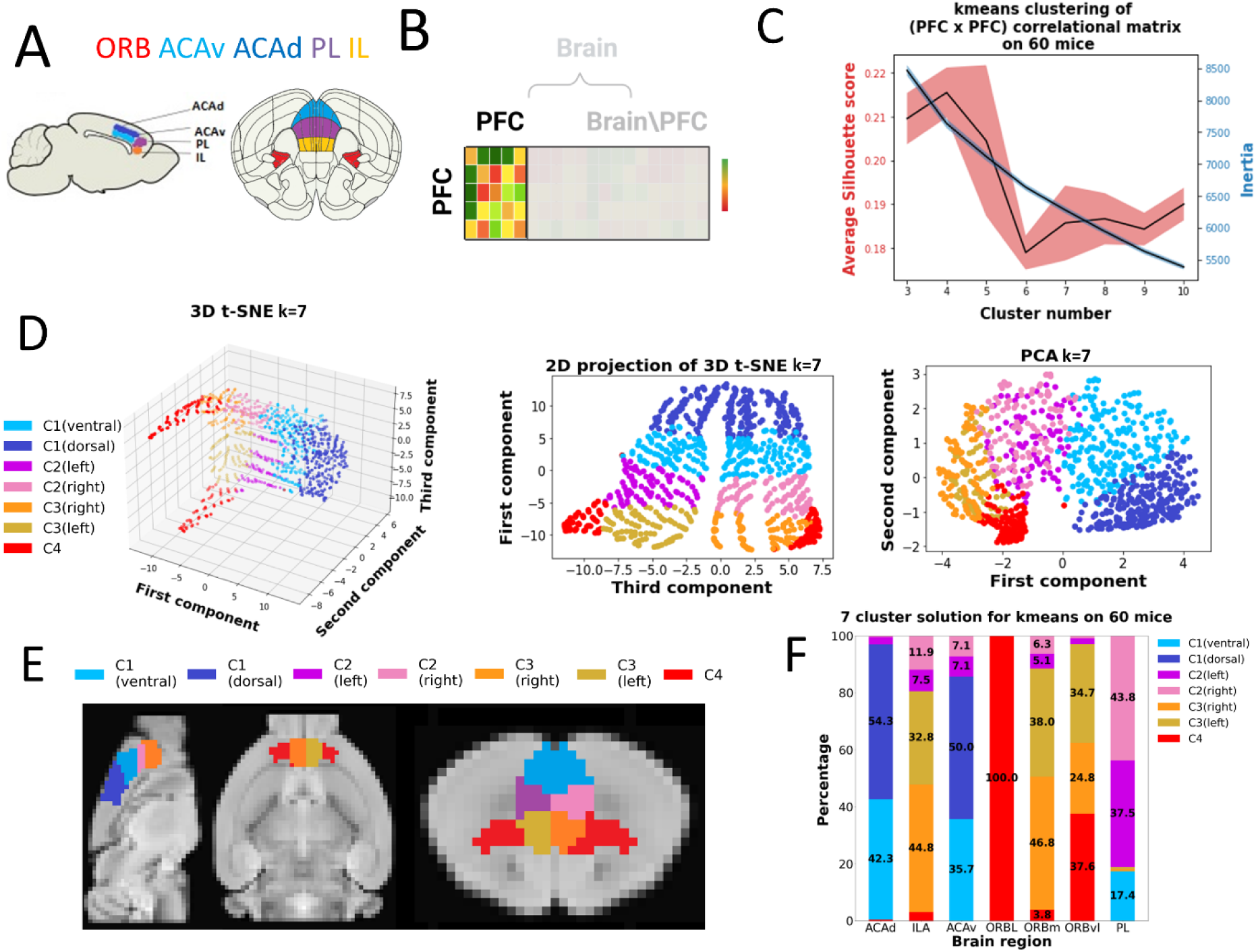
PFC clustering based on the local functional connectivity. **A:** Visualization of PFC and its subregions based on Allen Mouse Brain Common Coordinate Framework version 3 (CCFv3). **B:** Scheme of the correlation matrix used to derive PFC subdivisions via kmeans. **C:** Average silhouette and inertia scores for various cluster solutions using 60 randomly chosen mice. Black line represents the mean values across 200 iterations. Shaded area features the standard deviation. **D:** Low dimensional representation of the correlation matrix through 3D t-SNE and PCA for 7-cluster solution. Colors represent the clusters derived by k-means clustering on the complete (not-embedded) correlation matrix. **E:** Visualization of the anatomy of individual clusters along sagittal, transverse and coronal sections. **F:** Prevalence of individual clusters based on 7-cluster solution within anatomically defined (CCFv3) regions. Percentage values are depicted only for the regions with a contribution above 3%.

However, when looking at the short-range connectivity within PFC, the silhouette scores exhibited a second peak at 7-cluster solution, which was reproduced for various data sizes (Fig S4). This result indicates that FC-based parcellation based on short-range connection within the PFC could capture high-order anatomical subdivisions of the PFC. Importantly, cluster borders from 7-cluster solution were consistent across different data sizes and clustering methods (Fig S4 and Fig S5).

The newly identified clusters outline a further separation of FC along medial-lateral (C2(left)/(right), C3(left)/(right) and dorso-ventral axes: C1(dorsal)/C1(ventral). Importantly, all rostro-caudal splitting discussed for 4-cluster solution was preserved, additionally corroborating the stability of the 4-cluster solution. In particular, the combined contributions of C2(left)/(right), C3(left)/(right) and C1(dorsal)/C1(ventral) to the brain regions are comparable to the contributions from C2, C3 and C1 respectively (Fig 2F and Fig 1F). Also, the low dimensional representations of the local correlational matrix were similar to the ones produced at 4-cluster solution (Fig 2D and Fig 1D).

Splitting of C3 to C3(left)/(right) and C2 to C2(left)/(right) clusters along medial-lateral axis are presumably driven by the higher FC of connections within a hemisphere (e.g intrahemispheric) compared to the connections between left and right hemispheres (e.g interhemispheric). This lateral splitting is in accordance with the connectivity data from (Zingg et al., 2014) showing the intrahemispheric reciprocal pathway from SUBd to PL/IL via ACAv. As was suggested by Zingg et al., this pathway is responsible for conveying the spatial information from SUBd. Interestingly, we could previously identify the same pathway by analysing the combined contribution of C2 in ACA and PL for 4 cluster solution. Exposing left/right segregation for C2/C3 highlights the hemisphere specificity of the SUBd-PL/IL pathway, which is in accordance with existing tracing data (Zingg et al., 2014). Furthermore, because this 7 cluster FC-based splitting only arises when analyzing PFC-PFC connections, we propose that SUBd spatial information is largely processed inside intrahemispheric PL/IL networks. The identification of intrahemispheric local networks in the prefrontal cortex is consistent with Harris and colleagues’ hypothesis of the prefrontal subsystem, who revealed the dominance of intrahemispheric connections within the group of prefrontal areas, including ORB, ACA, PL, and IL using axonal tracing (Harris et al., 2019).

Segregation of C1 into C1(dorsal)/C1(ventral) is of particular interest since it provides an answer to a pending inconsistency in the demarcation of the cingulate cortex. In short, borders between subregions of cingulate cortex are drawn along rostro-caudal axis for most mammals (e.g. humans, non-human primates, rabbits), but not for mice and rats where the borders are defined perpendicularly to the same axis (Fig 2 A) (van Heukelum et al., 2020). As Fig 2E exhibits the FC-based clustering supports the demarcation along rostro-caudal axis and not along dorso-lateral, as it is done in widely used Allen (Wang et al., 2020) and Paxinos (Paxinos & Franklin, 2019) brain atlases.

### Using of FC-based clusters to discern the network effect of external perturbations

Targeted brain stimulation is being used increasingly to address neurological illnesses and improve human performance. To boost the effectiveness of these strategies, a key objective of these focused perturbations is to modulate activity within a certain network while having no influence on others (Frohlich & Riddle, 2021; Sullivan, Olsen, & Widge, 2021). In this study, we discovered that the mouse PFC can be defined by four major functional clusters, each characterised by a unique pattern of whole-brain connectivity (i.e. fingerprint) and inter-connectivity within the PFC. We argue that a perturbation approach would affect connectivity in downstream areas connected with the target region but will not further propagate to anatomically close but functionally unconnected PFC clusters.

In order to test this, we injected a ssAAV2-mCaMKIIα-hM3D(Gq)-mCherry-WPRE-hGHp(A) DREADD virus that targets excitatory, projecting neurons near the junction of the two C1 and C2 functional clusters (Fig 3A). During an fMRI session we then chemogenetically activated these cells and compared FC of each cluster before and after the activation to assess the effect and specificity of the chemogenetic perturbation (Fig 3B).

**Figure 3.**
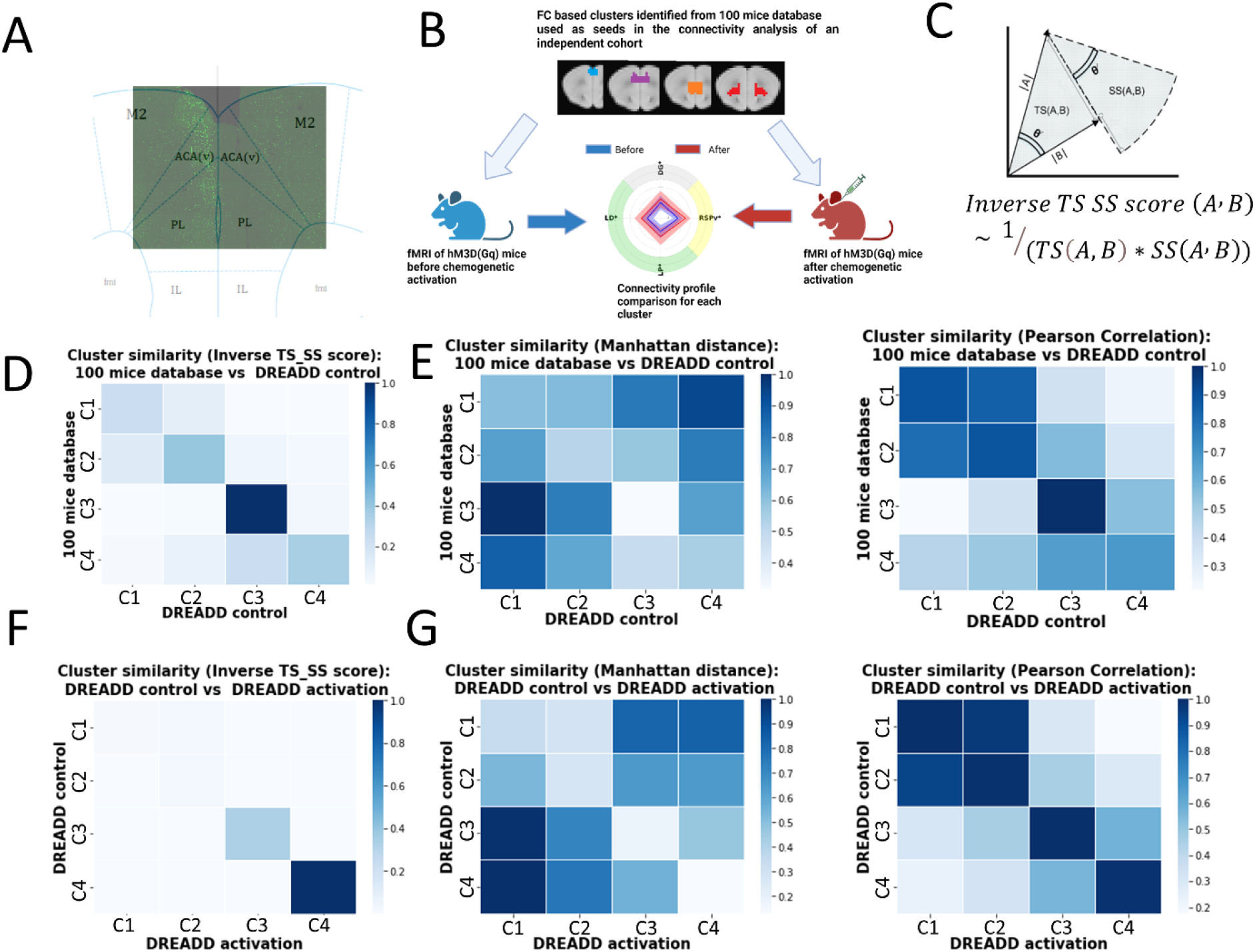
Cluster similarity characterization with geometry-based metric. **A:** cFOS activation induced by chemogenetic activation outlines the injection spot **B:** Experimental design **C:** Graphical representation of TS-SS similarity score **D-G:** Cluster similarity matrices computed using TS-SS, Manhattan and Pearson Correlation scores. All values are normalized by the division to the highest score computed using the chosen metric (see Methods for details).

Each fingerprint can be viewed as (1xN) dimensional vector, where N is the number of distinct brain areas, containing average z-scores for each brain area. We quantified the similarity between different fingerprints by using a combined metric based on Triangle’s Area Similarity (TS) and Sector’s Area Similarity (SS), named TS-SS score (Fig 3C) (Heidarian & Dinneen, 2016). In contrast to commonly used metrics such as: Pearson Correlation and Manhattan distances (Oduntan, Adeyanju, A.s, & Obe, 2018), that rely on either (i) an angle between compared vectors, therefore, not sensitive to magnitude changes (e.g. Pearson Correlation) or (ii) a difference between the vector’s magnitude in each dimension (e.g. Manhattan distance), TS-SS being a geometry-based metric uses both: the angle between the vectors and the difference in their amplitudes. We argue that the usage of both metrics: orientation and magnitude is of particular importance, since the connectivity fingerprints convey information about (i) strength of FC with a specific region (e.g. scale of a vector in a specific dimension) and (ii) the relative ratios of the FCs amplitude in various regions (e.g. orientation or an angle of the vector in a high dimensional space).

### Cluster solution computed on 100 mice database is reproduced by an independent dataset

Our first objective was to juxtapose connectivity fingerprints specific to clusters derived from the 100-mouse database with those from the experimental DREADD mice pre-chemogenetic activation (referred to as the "DREADD control group"). The inverse TS-SS scores, representing the similarity in cluster fingerprints, are illustrated in Fig 3D. Diagonal elements depict the similarity scores within the same clusters for the "DREADD control" and "database" groups, while off-diagonal elements indicate similarity scores across different clusters in these groups. Notably, the maximum values align with the diagonal, reinforcing the reproducibility of the identified four-cluster solution. Particularly, the fingerprints of C1/C2 and C3/C4 clusters exhibit significant overlaps, consistent with the previously discussed co-occurrence of clusters C3/C4 in the ORB area and C1/C2 in the ACA region (refer to Fig 1E). Importantly, these findings remained consistent across various similarity scores, as demonstrated in Fig 3E.

### Functional connectivity changes upon cluster-specific chemogenetic activation

Next, we sought to define the changes in whole-brain connectivity brought by our chemogenetic activation by contrasting cluster fingerprints before (i.e., DREADD control) and after the activation (i.e., DREADD activation).

The elevated similarity scores observed for C3 and C4 fingerprints, contrasted with lower scores for C1 and C2, suggest that the perturbation predominantly influenced C1 and C2 (Fig 3F). This aligns with our expectations, as the chosen anatomical and functional target was positioned at the interface between these clusters (Fig 3A). Interestingly, these changes were detectable using magnitude sensitive methods (Manhattan and TS-SS), which supports our argument that it is preferable to compare fingerprints using a metric that is sensitive to different geometric criteria (e.g. angle, magnitude) rather than restrict the choice to the metrics sensitive to one criteria (e.g. angle for Pearson Correlation) (Fig 3F and Fig 3G). To illustrate the primary impact on functional connectivities within C1 and C2, we conducted a comparison of averaged regional z-scores before and after ACA/PL activation (Fig 3B, Fig 4).

**Figure 4.**
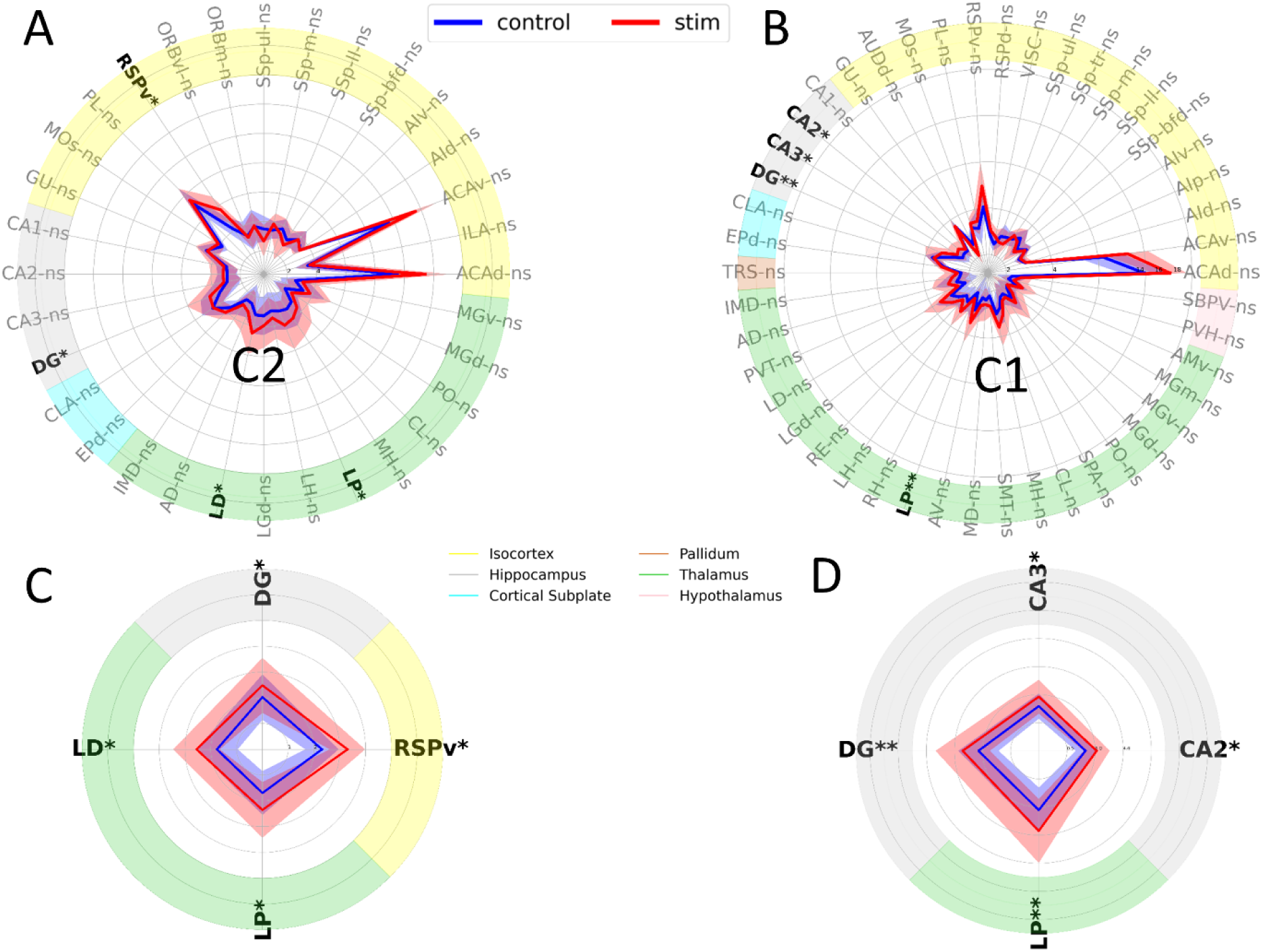
Fingerprints comparison between DREADD control and DREADD stimulation. Representation of the whole-brain activity for C2 and C1 clusters. Each value corresponds to an absolute average z-score across the corresponding group in the assigned anatomical regions. Only brain areas with absolute z-scores higher than 1.96 in at least one of the groups are shown. If for both groups at a specific area absolute z-scores are lower than 1.96, these areas are excluded from the comparisons to avoid false positives. Depending on the normality of the distributions either paired t-test or Wilcoxon signed rank test were used to estimate statistical significance. Resulted p-values were corrected for multiple comparisons using Bonferroni correction. See Table S1 for used abbreviations.

We found that the chemogenetic perturbation selectively affected (i) thalamic (LD and LP) areas, (ii) Retrosplenial cortex (RSP) and (iii) hippocampus (DG, C3, CA2), that all belong related to the well-established resting state network - Default Mode Network (DMN)(Perry & Mitchell, 2019; Whitesell et al., 2021; Zerbi et al., 2015) (Fig 4). No regional changes were found for clusters C3 and C4, which is consistent with their high similarity scores before and after the perturbation (Fig 3F, Fig S6).

Thus, by comparing similarities of clusters fingerprints we (i) additionally validated previously identified 4 cluster FC-based parcellation of PFC and (ii) identified the increase of FC only at clusters overlapping with the chemogenetically activated brain areas. It is important to note that the perturbation’s impacts did not extend to downstream neighbouring anatomical regions and off-target bordering clusters. We contend that these findings provide additional support for the robust network-level segregation of the mouse mPFC and demonstrate how FC-based parcellation may be used to identify the regional specificity an external perturbation.

## Discussion

### Functional connectivity-based parcellation of the mouse prefrontal cortex

The PFC is a critical brain region that aids in sensory integration, the processing of information linked to motivation and rewards (Padoa-Schioppa & Assad, 2006)and the application of reward-related strategies and behaviors (Grueschow, Polania, Hare, & Ruff, 2015; Hare, Camerer, & Rangel, 2009). Functional magnetic resonance imaging (fMRI) studies revealed an elevated metabolic activity in ventral medial portion of PFC (vmPFC) in patients with major depression disorder (MDD) (Koenigs & Grafman, 2009; Ubl et al., 2015), leading to the usage of vmPFC as a therapeutic target for Deep Brain Stimulation (DBS) in MDD(Mayberg et al., 2005; Moreines, McClintock, Kelley, Holtzheimer, & Mayberg, 2014; Sullivan et al., 2021). Notably, psychiatric conditions (e.g. MDD) are thought to result from malfunctions of distributed brain networks rather than an aberrant activity of a single isolated region. Therapeutically this means that rather than restoring an abnormal activity of a single brain area, it is more effective to adjust the applied stimulation to normalize deviant network functioning (Sullivan et al., 2021). For MDD, it has been suggested that selecting a target for DBS based on individual aberrant connectivity would yield optimal results (Zhu et al., 2021). Following these considerations for a network-specific targeting approach, we posit that, similar to the human-based studies described above, rodent research would benefit from directing attention toward specific functional networks rather than adhering to a predefined anatomical area. In the present study, we employed functional connectivity (FC)-based parcellation of the mouse prefrontal cortex (PFC) and identified four stable and reproducible co-activation clusters within the PFC. Through the analysis of overlap between the revealed clusters and anatomically defined regions, we demonstrated that the cluster locations align with the established anatomical segregation of the PFC along the dorso-ventral axis into the anterior cingulate area (ACA), prelimbic cortex (PL), infralimbic cortex (IL), and orbital cortex (ORB) areas (Le Merre et al., 2021). Additionally, we demonstrated that further subdivision of PL and IL is dominated by short-range (within PFC) intrahemispheric connections similarly as was previously suggested by tracing studies (Harris et al., 2019; Zingg et al., 2014). At the same time, functional segregation of ACA occurred along rostro-caudal axis, matched anatomical subdivision used for humans and opposed the dorso-ventral demarcation commonly used in rodent atlases, potentially providing an answer to a long-lasting inconsistency in cross-species ACA segregation (van Heukelum et al., 2020).

### Non-overlapping whole-brain functional networks linked to PFC clusters

To examine the functional boundaries of our clusters, we employed these functional areas as targets for a chemogenetic neuronal excitation protocol. We observed that the impact of the applied stimulation on whole-brain functional networks is linked with the targeted functional area as suggested by previous work (Mandino et al., 2019; Markicevic et al., 2020).

Specifically, our chemogenetic stimulation resulted in FC alterations within targeted brain areas, while neighboring clusters showed no detectable network changes, highlighting significant functional segregation in the PFC. Notably, brain regions exhibiting the most significant FC changes within perturbed networks were distant, including the hippocampus, thalamus, and retrosplenial cortex, aligning with the previously identified DMN. Surprisingly, FC changes in nearby anatomical regions (ACA, PL, IL, ORB) were not statistically significant. This intriguing outcome suggests that targeting the area with aberrant connectivity may be more efficiently achieved by perturbing the associated network rather than directly focusing on the specific anatomical region.

### Conclusion

In this investigation, we employed correlation functional connectivity metrics to successfully parcellate the mouse PFC, revealing stable and reproducible clusters. Utilizing these clusters as targets for chemogenetic activation, our findings demonstrated that the activation effects were confined to the target clusters, exhibiting no influence on surrounding anatomical areas or neighboring off-target clusters. The observed localized effects of external perturbation underscore the remarkable network-level segregation within the mouse mPFC. This study contributes valuable insights into the network-specific organization of the mouse PFC, highlighting its potential implications for targeted interventions.

## Methods

### Data processing

All processing steps were performed using FSL and custom-made scripts at Python 3.7 using SCIPY, SKLEARN, NIBABEL, NUMPY, NILEARN toolboxes, as well as custom-build library for TS-SS similarity computation (Heidarian & Dinneen, 2016). Illustrations were created with Biorender.com.

The principal steps of the data processing are summarized in Fig 5.

**Figure 5.**
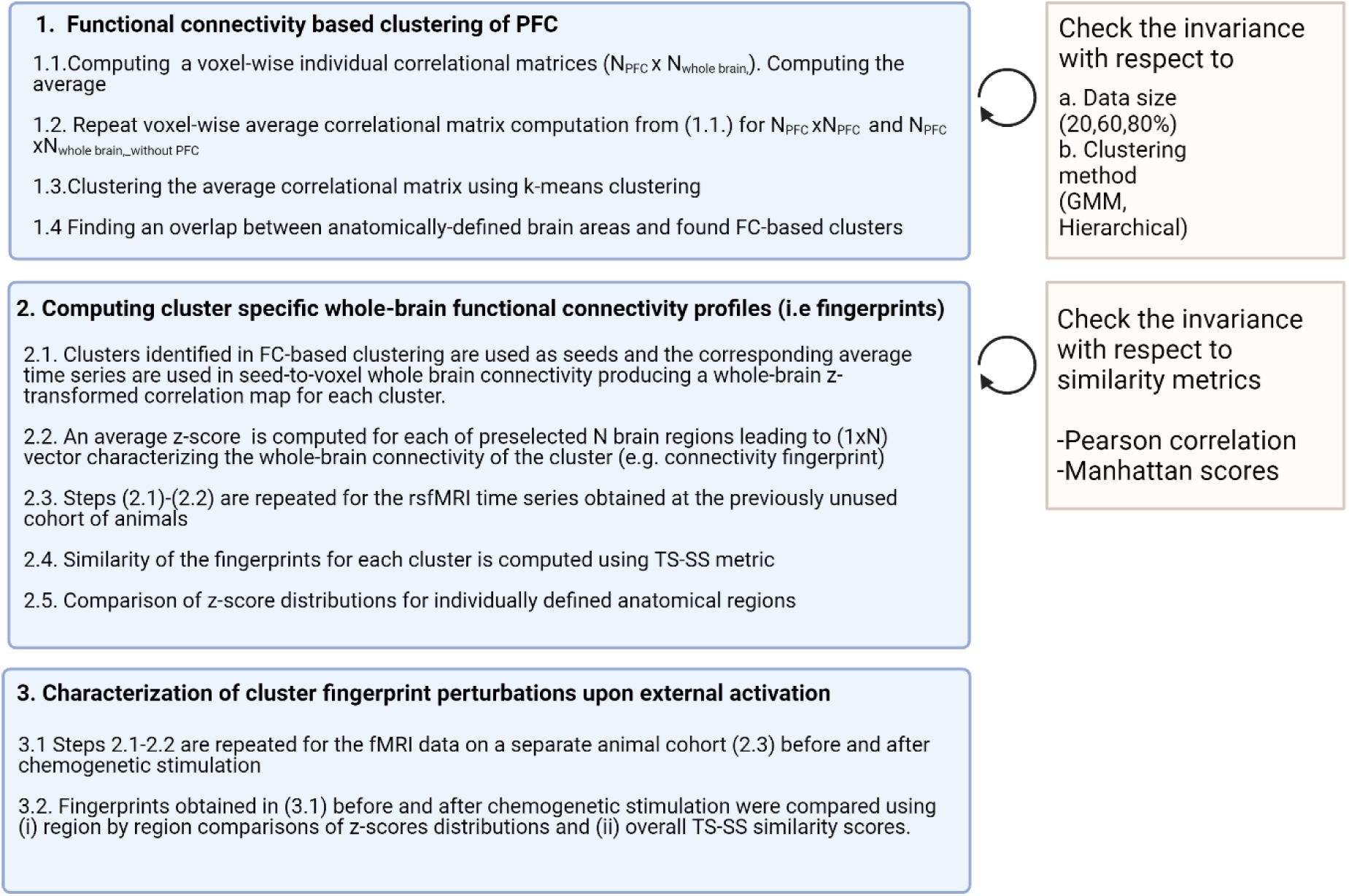
Data processing steps. Chart summarize the main steps done for the data processing that can be divided into 3 categories (“blue boxes”) and the analysis to insure the invariance of the obtained results (“gold boxes”). Details of each section are provided in the corresponding subchapters of Data processing paragraph.

### Functional connectivity-based clustering of PFC

Firstly, BOLD timeseries were extracted using NIBABEL toolbox and depending on the type of the computed correlational matrix (e.g [PFC x BRAIN], [PFC x PFC] or [PFC x BRAIN without PFC]) the corresponding masks were used to select the voxels of interest. Specifically, as BRAIN we considered the combination of the following macroscopic regions according to Allen Mouse Brain Common Coordinate Framework version 3 (CCFv3)(Wang et al., 2020): isocortex, striatum, thalamus, hypothalamus, hippocampus (without Entorhinal area, Presubiculum, Postsubiculum and Parasubiculum), cortical subplate and pallidum. List of the analyzed 122 brain areas is at Table S1. Importantly, similar to (Le Merre et al., 2021) we did not include Anterior Insular cortex (AI) in PFC. MOp was, also, excluded from PFC because we wanted to concentrate on high-order processing occurring in PFC, akin to (Harris et al., 2019).

The ConnectivityMeasure function from the NILEARN toolbox was used to further generate correlational matrices for each animal using masked timeseries. The average correlational matrix was calculated using the appropriate number of matrices, depending on the selected data size (20,60,80, or 100 mice). The computed average correlational matrix was clustered using SKLEARN toolbox applying either Kmeans clustering, Agglomerative clustering with Ward linkage (i.e. Hierarchical clustering) or Gaussian Mixture Models with full covariances and “kmeans” as initial parameters (Fig 1-2, Fig S1-S5). The optimal cluster solution was chosen by identifying a maximum peak at the average silhouette score plot (Rousseeuw, 1987). Additionally, for Kmeans clustering the cluster choice was verified by finding an “elbow” point (Thorndike, 1953) at the Inertia plot. To check the reproducibility of the identified cluster solutions at different sample sizes and to be independent from the exact sample choice we performed random data resampling without repetitions (N=200) at various group sizes (20,60,80%) and computed distributions of Average Silhouette Scores and Inertia.

Finally, we overlaid the coordinates of the voxels assigned to each cluster with the known coordinates of anatomically defined regions to determine the relative presence of the clusters in each region of PFC.

### Computing cluster specific fingerprints

Identified FC-based clusters are further used as seeds for seed-to-voxel whole brain connectivity analysis using FSL (Smith et al., 2004) as previously described in (Zerbi et al., 2015). For each seed (i) an average time course across the seed voxels was computed and used to calculate the Pearson correlation between the seed and the voxels outside of the seed. The resulted correlation map was z - transformed.

An average z-score across the voxels was calculated for all previously listed 122 brain regions, generating a vector (1x122) that represents the cluster’s whole-brain connectivity (i.e “fingerprint”) for each animal.

To check the reproducibility of the identified cluster solution the cluster fingerprints were computed for the previously unused experimental group (n=13) (“DREADD control”) (scan time 0-15 min, see MRI setup for details). Three complimentary similarity techniques were used to assess the similarities of the fingerprints: Pearson correlation (Kornbrot, 2005), Manhattan Distance (Krause, 1973) and Triangle Area Similarity (TS) -Sector Area Similarity (SS) (i.e. TS-SS score) (Heidarian & Dinneen, 2016) (Fig 3D-G)

For two fingerprints x and y of length N theses similarities were computed using the formulas (1 - 8):

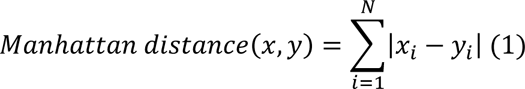

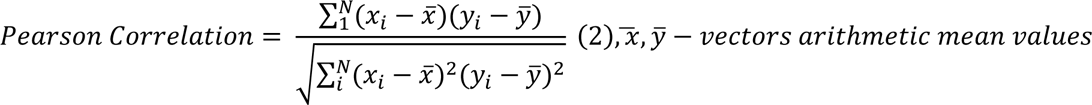

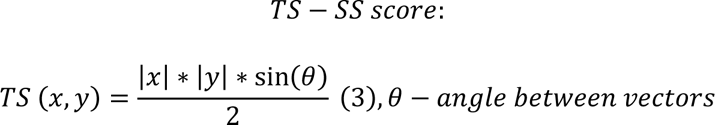

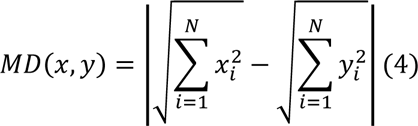

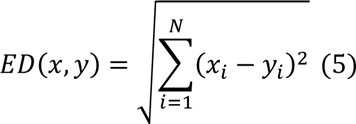

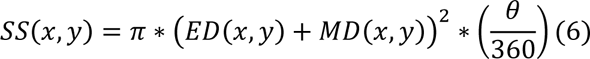

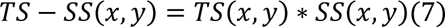

For the similarity comparisons an average fingerprint across all animals was computed for each cluster. To compute similarity matrices in Fig 3 similarity scores for each method were normalized according to formula (8).

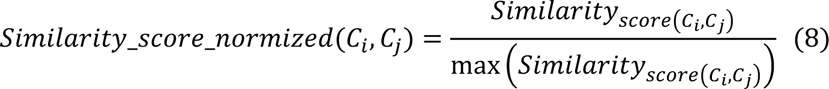

where C_i_ and C_j_ are clusters I and j (i=1..4, j=1..4). Similarity_score(C_i_, C_j_) corresponds to the similarity score computed using formula (1), (2) or (7).

### Characterization of cluster fingerprint perturbations upon external activation

To assess the effect of the external perturbation on each cluster fingerprint the fingerprints were computed for previously mentioned DREADD control group after the DREADD activation – “DREADD stimulation” (scan time 23-38 minutes see MRI setup for details) (Fig 3 F-G).

To pinpoint areas conveying most of the dissimilarities between the clusters we compared regional distributions of z-scores using paired t-test or Wilcoxon signed rank test and correction for multiple comparisons using Bonferroni correction(Bonferroni, 1936) (Fig 4 and Fig S6). To increase the robustness of the regional z-score comparison, we only included the regions for which the distribution of voxels absolute z-scores is statistically significantly higher than 1.96 for at least one of the compared fingerprints.

## Experimental procedures

### 100 mice database

Previously acquired rsfMRI scans (Zerbi et al., 2021) were used. Anesthesia and MRI setup parameters match the corresponding sections below: Anesthesia for MRI experiment and MRI setup, with two adjustments (i) no clozapine injection was performed and (ii) scanning time was 60 min.

### Animals

All animal experiments were performed under license from the Zürich Cantonal veterinary office following Swiss federal guidelines for the use of animals in research. For the experiments C57BL/6J mice (Charles River Laboratories, Germany) were used. Animals were housed in IVC cages in a group of five with 12-hour light/dark cycle, ad libitum food in temperature and humidity-controlled ETH Zürich facility (EPIC).

### Anesthesia

Anesthesia protocols were carried as previously described in (Markicevic et al., 2020)

### Anesthesia for MRI experiment

All mice were initially anesthetized with isoflurane (in a 1:4 O2: air mixture as the carrier gas): 4% for induction, 2% for endotracheal intubation and during set-up on the animal bed. After the animal was placed in the MRI bed, it was connected to a small animal ventilator and mechanically ventilated at a rate of 80 breaths/min, with isoflurane set to 2%, with a respiration cycle of 25% inhalation, 75% exhalation, and an inspiration volume of 1.8 ml/min.

The mouse head was immobilized using an incisor tooth bar and non-rupture ear bars. Body temperature was measured using a rectal temperature probe and maintained at 36.5± 0.5°C by means of a warm-water circuit integrated into the animal holder (Bruker Biospin GmbH, Ettlingen, Germany). Following animal preparation, the tail vein was cannulated for intravenous (i.v.) administration of a bolus of 0.05 mg/kg medetomidine (Domitor, medetomidine hydrochloride; Pfizer Pharmaceuticals, Sandwich, UK) and pancuronium (0.25mg/kg), isoflurane flow was lowered to 1%. After 5 minutes, continuous infusion of medetomidine (0.1 mg/kg/h) and pancuronium (0.25 mg/kg/h) was started. Isoflurane was lowered to 0.5%. An anesthesia/pancuronium solution was freshly prepared every day, by mixing saline, stock solution of medetomine 1 mg/ml, and stock pancuronium 1 mg/ml. Bolus volume and infusion rate were adjusted to the animal body weight to match the desired dose.

### Anesthesia for terminal perfusion

For brain collection followed by histology, terminal cardiac perfusion was carried out under deep anesthesia using i.p. injections of a combination of ketamin (100 mg/kg)/ xylazin (10 mg/kg)/ acepromacinmaleat (2 mg/kg). The invasive procedure was not started before pedal withdrawal reflex has disappeared (additional dosing until reached).

### Anesthesia for stereotaxic surgery

The mouse was initially anesthetized with 4% isoflurane (in a 1:4 O2: air mixture as the carrier gas) and its head be placed in a stereotaxic head frame between non-rupture ear bars on a heating pad. The animal was further intramuscular injected with a mixture of fentanyl (0.05 mg/kg), midazolam (5 mg/kg) and medetomidine (0.5 mg/kg) for the anasthesia during the stereotaxic surgery. At the time of the anesthesia the animal’s body temperature was measured with a rectal probe (Harvard Apparatus, United States) and the respiration rate was monitored by the surgeon.

Upon the finish of the surgery antagonist-antidote mixture was injected into the mouse consisting of temgesic (0.2mg/kg), annexate (0.5mg/kg) and antisedan (2.5mg/kg). During the first 3 days after the surgery analgesic (Ketoprofen (10 mg/kg)) was administered on the daily basis to ameliorate potential pain during the recovery period (at least 2 weeks).

### Stereotaxic surgery for the DREADD injection

Stereotaxic surgery was performed as previously described in (Markicevic et al., 2020). The animal was anesthetized as previously described in Anesthesia for stereotaxic surgery until it no longer responded to pedal withdrawal reflex test. Surgery began only after the disappearance of the reflex. The scalp was cleansed and the hair on the scalp was removed either using an electric trimmer followed by an application of a depilatory cream.

After calibrating the automated coordinate system, the mouse was placed on a heating pad (Harvard Apparatus, United States) to support body temperature maintenance and then on a stereotactic frame. Non-rupture ear bars and the mouthpiece of the stereotactic frame were used to secure the mouse head. Eye ointment was applied to prevent eyes from drying during the surgery. Surgical blade was applied for vertical incisions.

A Hamilton syringe was used for injecting 200nl of ssAAV8-mCaMKIIα-hM3D(Gq)-mCherry-WPRE-hGHp(A) (serotype 2, VVF Zürich) targeting AP 1.9; ML -0.35 and DV at -2.8/-2. In the pilot study the target coordinates were determined, and the appropriate viruses (AAV) were be injected. Once the injection was performed, self-absorbing wound sutures were applied to close the wound.

### MRI Setup

MRI sessions were performed as previously described in (Markicevic et al., 2020; Zerbi et al., 2021). Using 7-T Bruker BioSpec scanner equipped with a PharmaScan magnet with receive-only cryogenic coil (Bruker BioSpin AG, Fällanden, Switzerland). Animals were anesthetized as described in anesthesia for fMRI experiment. Adjustment procedures included wobbling reference power and shim gradients adjustments using Paravision v6.0. Echo Planar Imaging (EPI) sequence was used for BOLD fMRI acquisition with the following parameters: repetition time (TR) = 1 s, echo time (TE) = 15 ms, flip angle = 60°, matrix size = 90 × 50, in-plane resolution = 0.2 × 0.2 mm2, number of slices = 20, slice thickness = 0.4 mm, and 2280 volumes for a total scan of 38 min. After 15 minutes of EPI acquisition a bolus of 10 µg/kg clozapine was intravenously injected to activate DREADDs. We adjusted the stimulation condition to begin at 23 minutes after the scan started, accounting for 8 minutes for the DREADD’s progressive activation based on our prior findings (Zerbi et al., 2019).

### cFOS staining

cFos staining was performed as previously described in (Zerbi et al., 2019). At the last scanning day after 90 minutes from the injection timepoint terminal anesthesia was applied and the animal was transcardially perfused with 4% paraformaldehyde (PFA, pH = 7.4).

The brain was blocked in cold 4% paraformaldehyde for 2h, following the rinsing with PBS and overnight storage in a sucrose solution (30% sucrose in PBS). For the long term storage the tissue was frozen in the mounting medium (Tissue-Tek O.C.T Compound, Sakura Finetek Europe B.V., Netherlands). For immunohistochemistry brain was sectioned coronally using a cryostat (Leica CM3050 S, Leica Biosystems Nussloch GmbH) into 40 μm thick sections. The sections were intubated with rabbit anti-cFOS (226 003, Synaptic Systems, 1:5000) primary antibody solution mixed with 0.2% Triton X-100, and 2% normal goat serum in PBS over 2 nights. After washing with PBS, the sections were transferred in goat anti-rabbit Alexa Fluor 488 (A-11008, Thermo Fisher Scientific, 1:500) secondary antibody solution.

Microscopy images were acquired in a confocal laser-scanning microscope (CLSM 880, Carl Zeiss AG, Germany) with pinhole aperture of 1.0 Airy Unit and image size 1024x1024 pixels.

## Supplementary materials

**Table S1.**
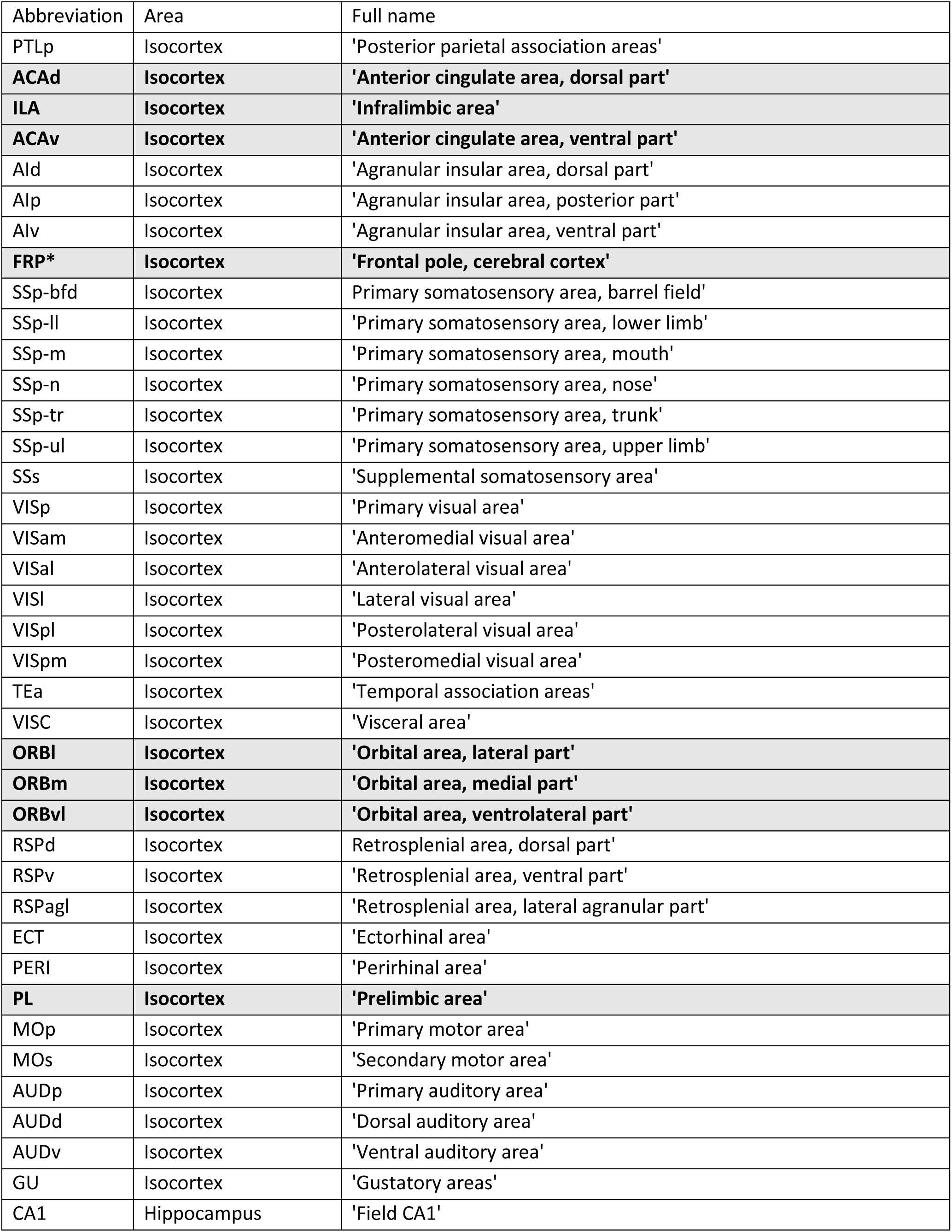
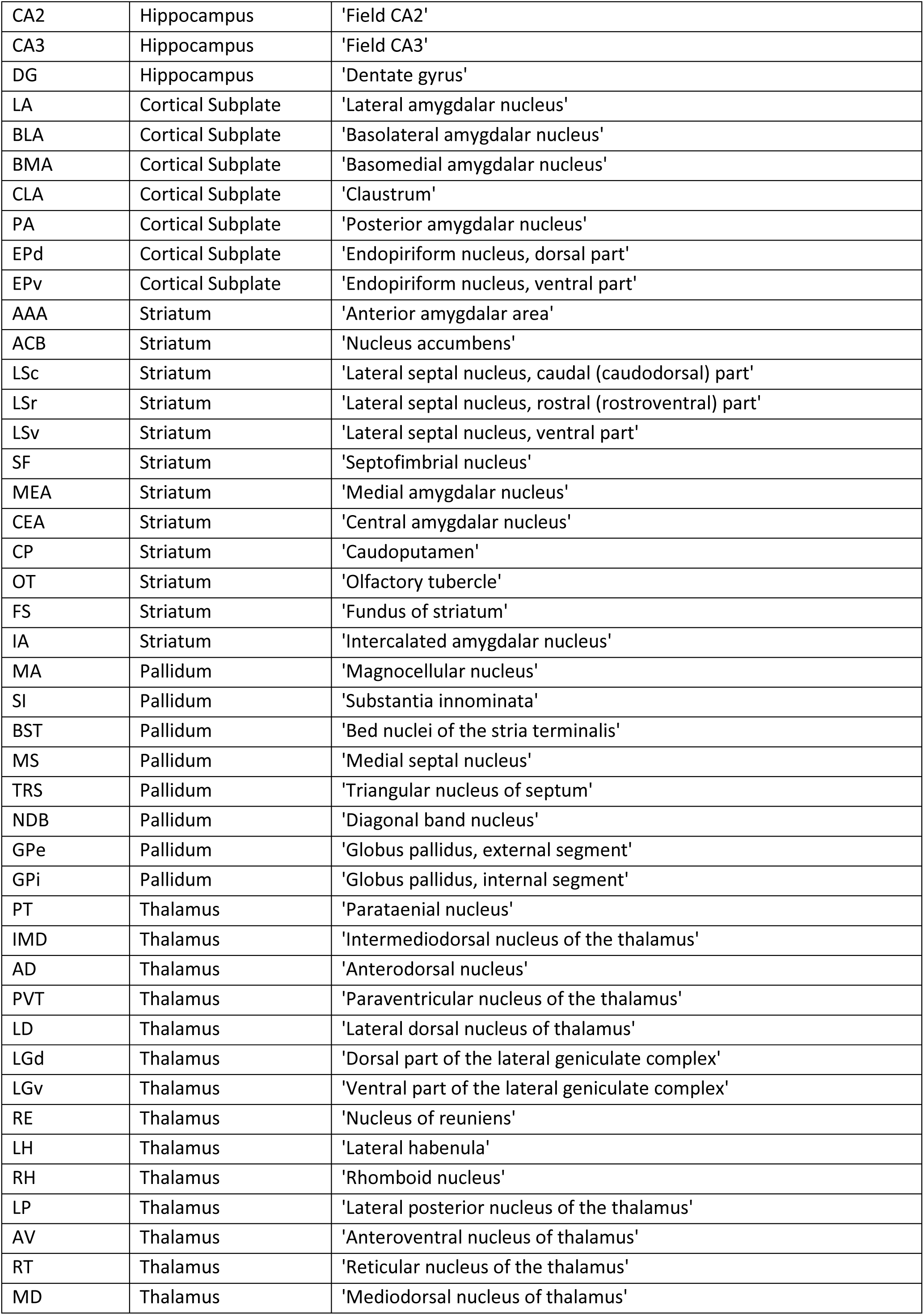
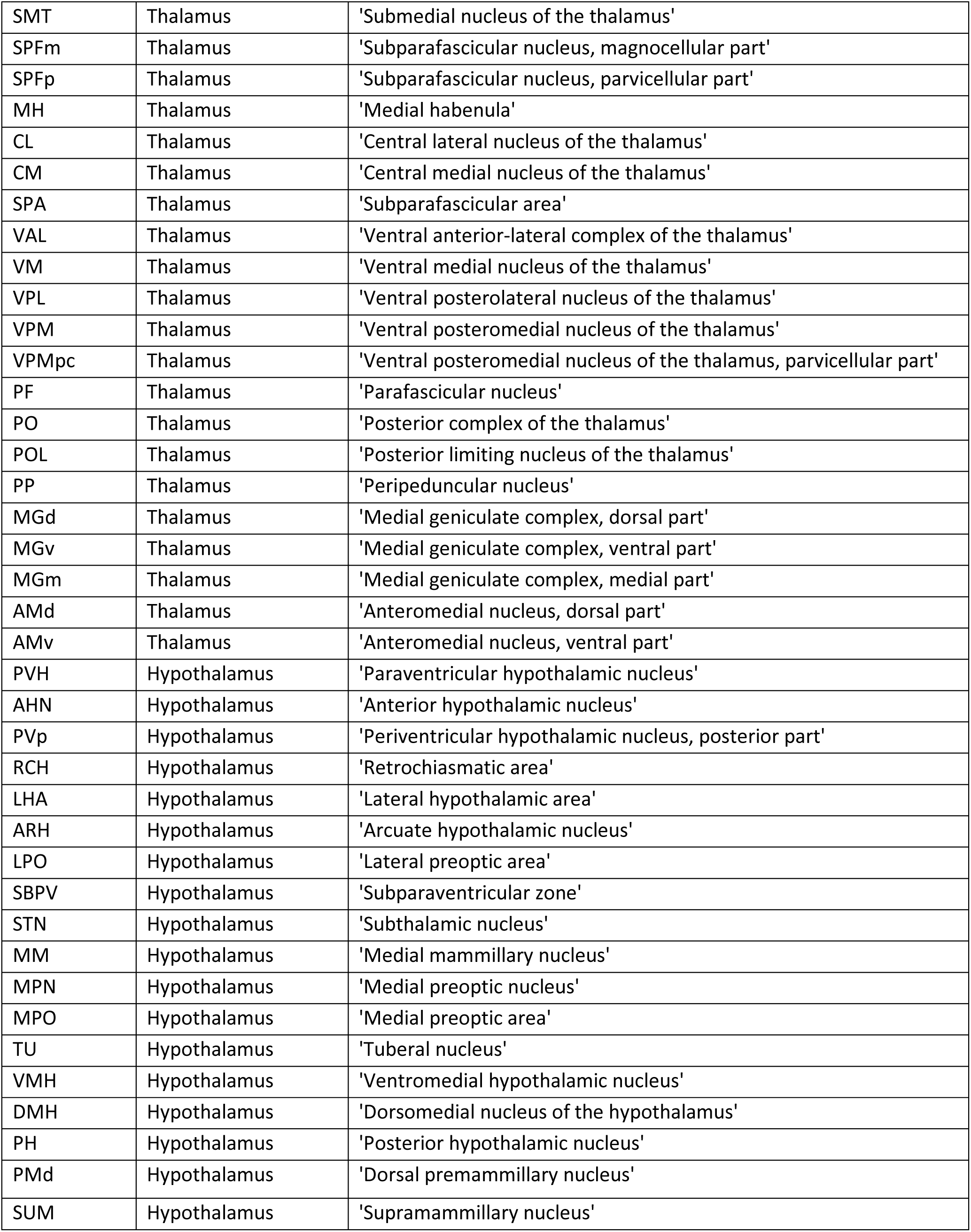
List of 122 analyzed anatomical areas named according to Allen Mouse Brain Common Coordinate Framework version 3 (CCFv3). Areas of Isocortex assigned to Prefrontal Cortex are additionally highlighted. *Most of FRP was outside of the scanned volume and, therefore, no cluster prevalence analysis (Fig 1F and Fig 2F) was conducted for this area. Signal from scanned FRP voxels was included in FC analysis (Fig 3-4)

**Figure S1.**
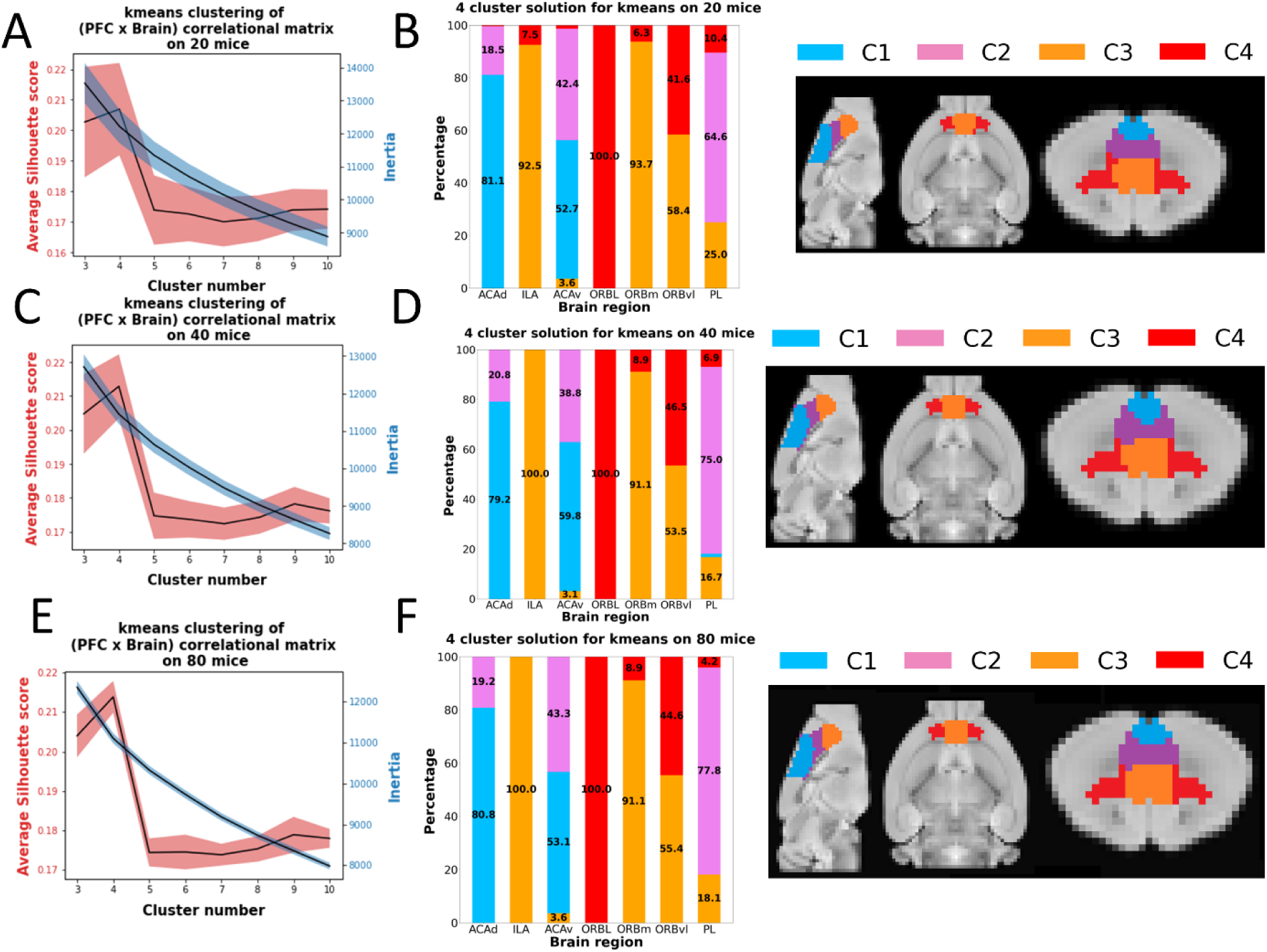
Whole-brain FC-based PFC clustering using different data sizes. **A,C,E:** Average silhouette and inertia scores for various cluster solutions using 20,40 or 80 randomly chosen mice. Black line represents the mean values across 200 iterations. Shaded area features the standard deviation. **B,D,F:** Prevalence of individual clusters based on 4-cluster solution within anatomically defined (based on CCFv3) regions. Percentage values are depicted only for the regions with a contribution above 3% (left). Visualization of the anatomy of individual clusters along sagittal, transverse and coronal sections (right).

**Figure S2. Whole-brain FC-based PFC clustering (k=4) using different clustering techniques.**

Visualization of the anatomy of individual clusters along sagittal, transverse and coronal sections for 4-cluster solution derived through kmeans, gmm and hierarchical clustering using complete 100 mice data set (left). Prevalence of individual clusters based on 4-cluster solution within anatomically defined (based on CCFv3) regions. Percentage values are depicted only for the regions with a contribution above 3% (right).

**Figure S3.**
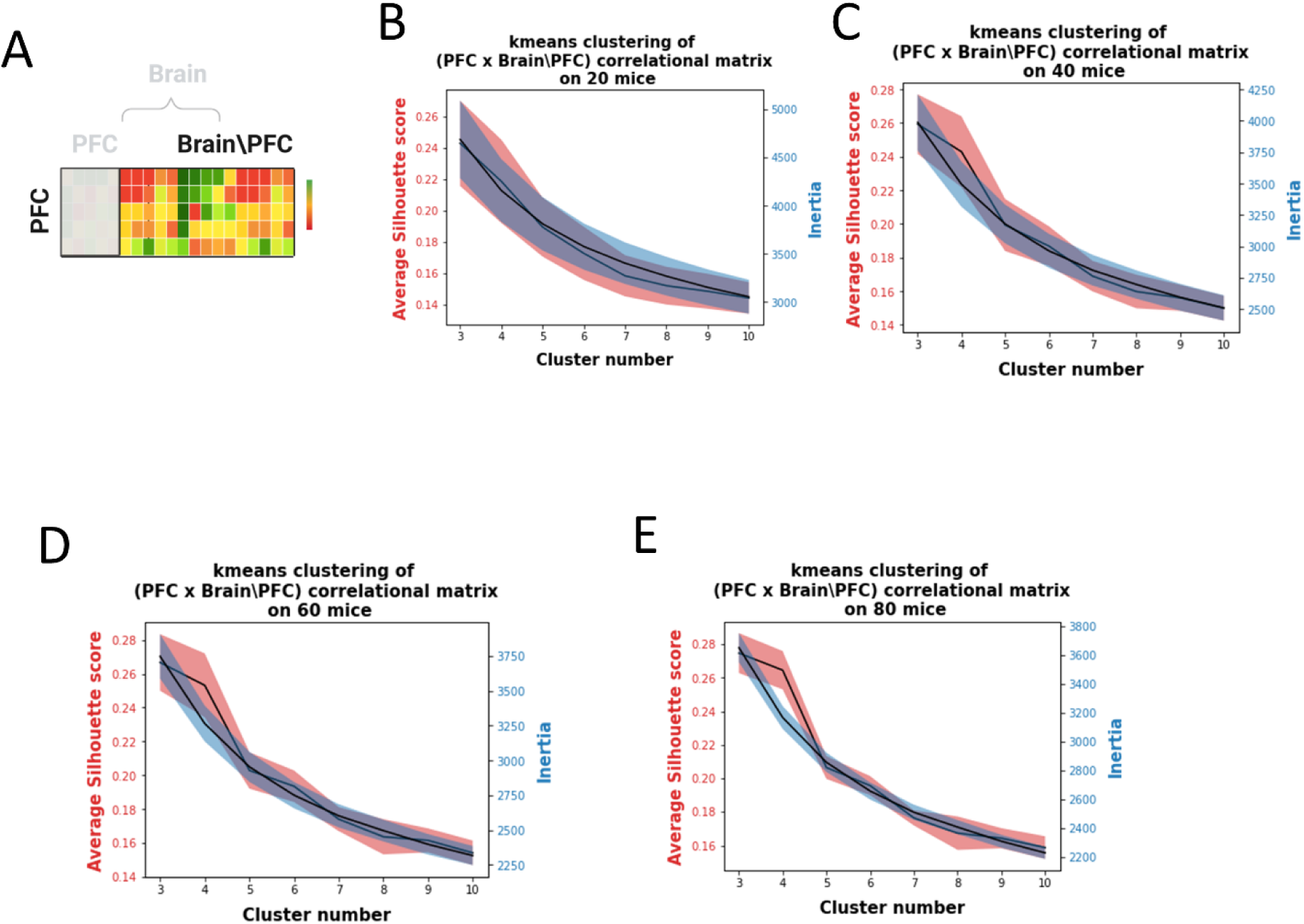
Diffusion FC-based PFC clustering using different data sizes. **A:** Scheme of the correlation matrix used to derive PFC subdivisions via kmeans. **B-E:** Average silhouette and inertia scores for various cluster solutions using 20,40,60 or 80 randomly chosen mice. Black line represents the mean values across 200 iterations. Shaded area features the standard deviation.

**Figure S4.**
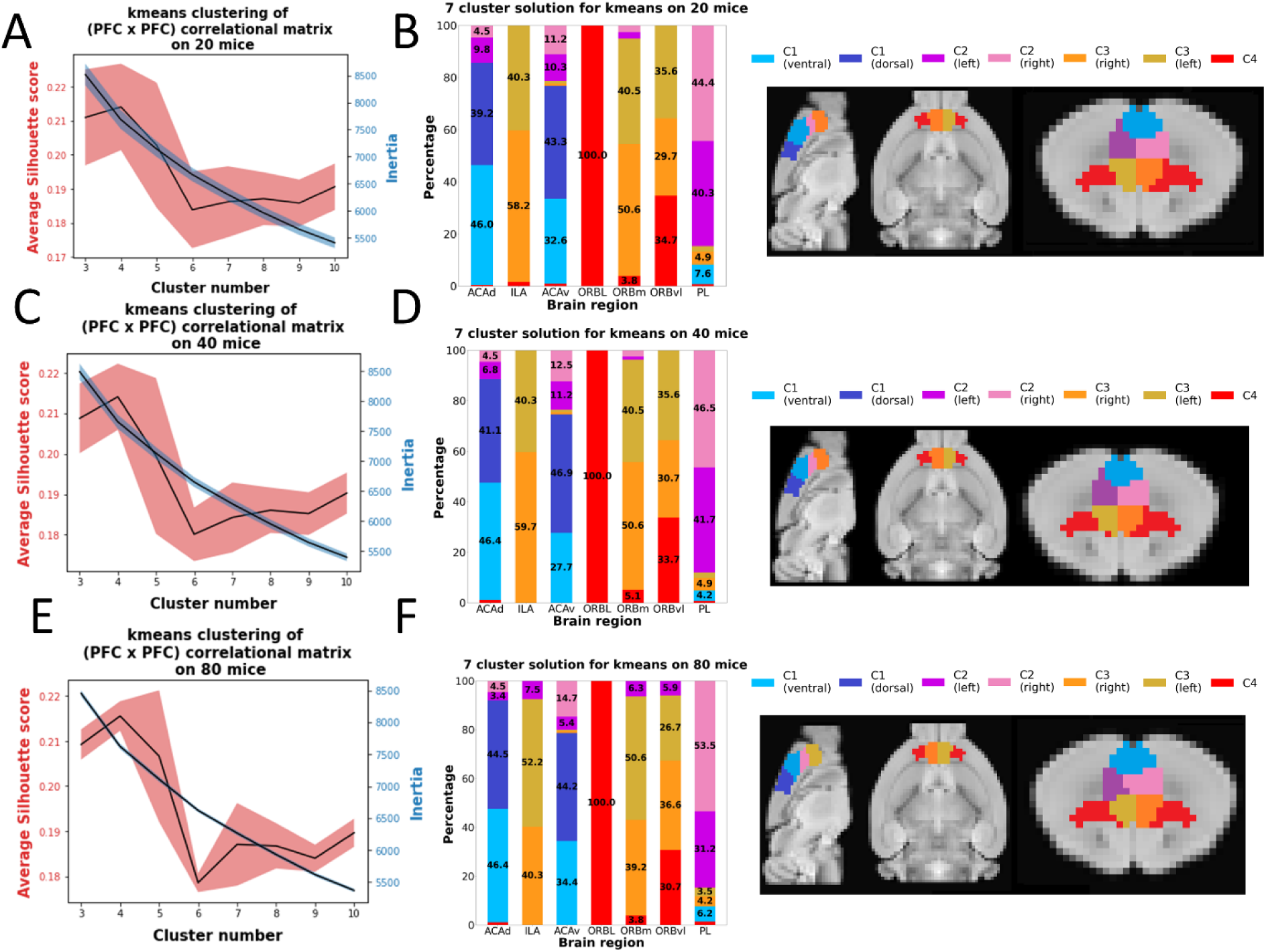
Local FC-based PFC clustering using different data sizes. **A,C,E:** Average silhouette and inertia scores for various cluster solutions using 20,40 or 80 randomly chosen mice. Black line represents the mean values across 200 iterations. Shaded area features the standard deviation. **B,D,F:** Prevalence of individual clusters based on 7-cluster solution within anatomically defined (based on CCFv3) regions. Percentage values are depicted only for the regions with a contribution above 3% (left). Visualization of the anatomy of individual clusters along sagittal, transverse and coronal sections (right).

**Figure S5.**
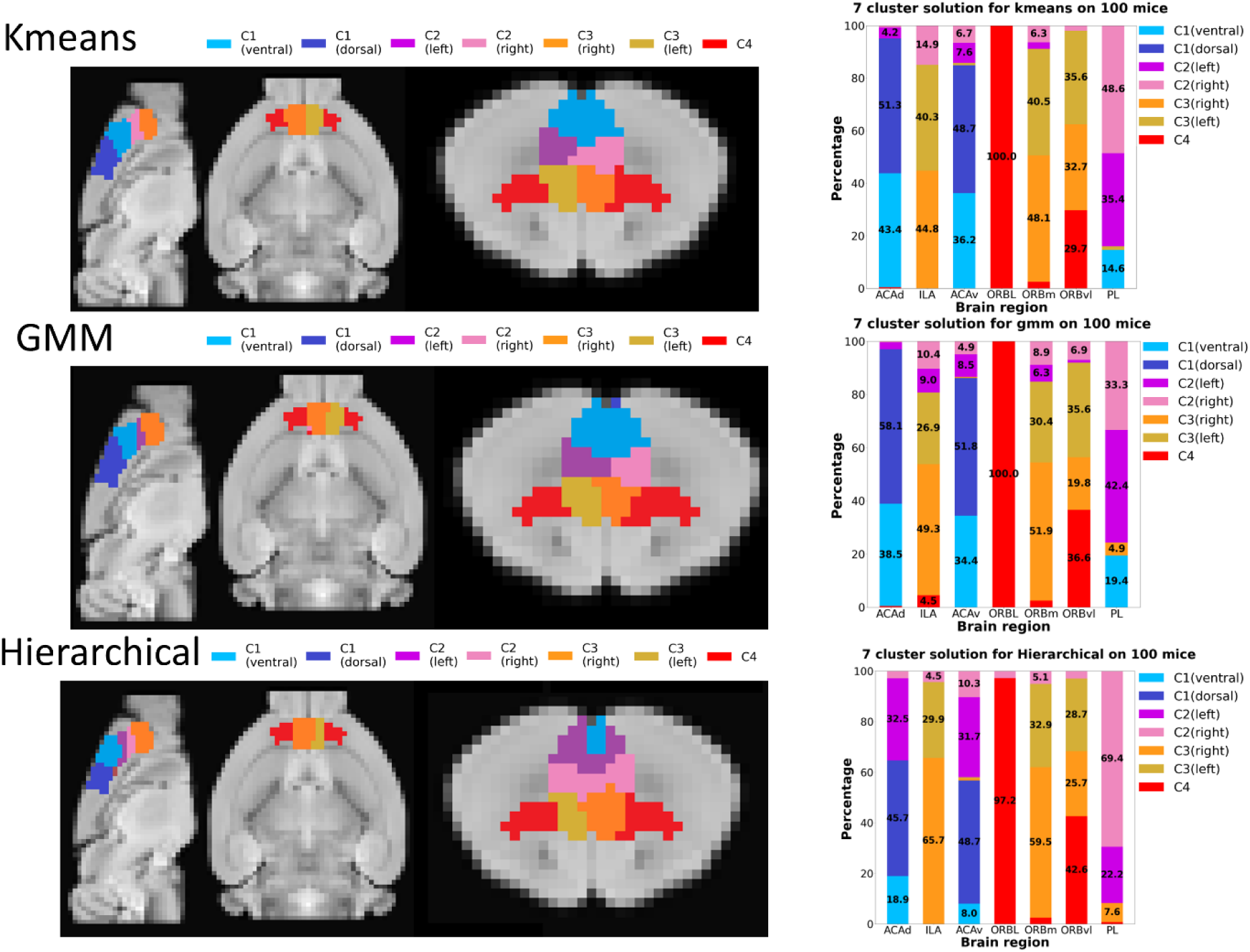
Whole-brain FC-based PFC clustering (k=7) using different clustering techniques. Visualization of the anatomy of individual clusters along sagittal, transverse and coronal sections for 7-cluster solution derived through kmeans, gmm and hierarchical clustering using complete 100 mice data set (left). Prevalence of individual clusters based on 7-cluster solution within anatomically defined (based on CCFv3) regions. Percentage values are depicted only for the regions with a contribution above 3% (right).

**Figure S6.**
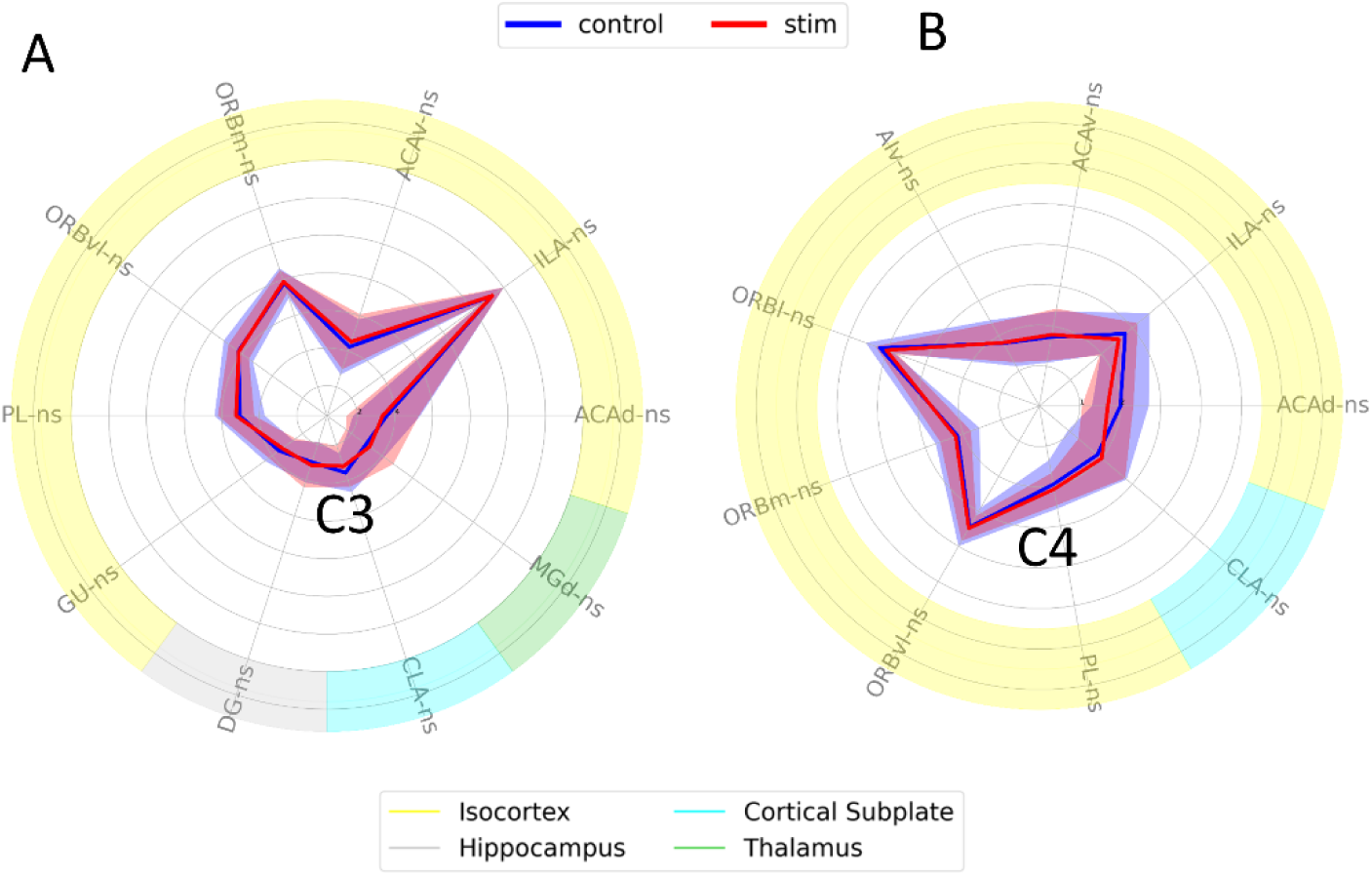
Fingerprints comparison between DREADD control and DREADD stimulation. Representation of the whole-brain activity for C3 and C4 clusters. Each value corresponds to an absolute average z-score across the corresponding group in the assigned anatomical regions. Only brain areas with absolute z-scores higher than 1.96 in at least one of the groups are shown. If for both groups at a specific area absolute z-scores are lower than 1.96, these areas are excluded from the comparisons to avoid false positives. Depending on the normality of the distributions either paired t-test or Wilcoxon signed rank test were used to estimate statistical significance. Resulted p-values were corrected for multiple comparisons using Bonferroni correction. See Table S1 for used abbreviations.

## Contributions

VZ, NW and RP acquired funding. VZ and IS conceived the project. CG performed chemo-fMRI experiments and followup histologies. MM, CG and VZ performed rsfMRI experiments. IS performed parcellation data analysis and cluster similiraty comparisons. IS and VZ wrote the manuscript and integrated corrections from all other authors. JG and RP reviewed the manuscript. All authors contributed to the article and approved the submitted version.

## Acknowledgements

This work was supported by an ETH Grant (ETH-25 18-2) to R.P. and V.Z., by a Swiss National Science Foundation (SNSF) ECCELLENZA (PCEFP3_203005) to V.Z., and by a European Research Council (ERC) starting grant (ENTRAINER) to R.P. This project has received funding from the European Research Council (ERC) under the European Union’s Horizon 2020 research and innovation program (grant agreement No. 758604).

We thank Markus Rudin for the important discussions

